# Bbl1 Allele of the Receptor Kinase Subfamily VII PBL38 Is Required for Elongated Root Hairs in Arabidopsis

**DOI:** 10.1101/2024.06.04.597371

**Authors:** Bruce D. Kohorn

**Affiliations:** Biology Department, Bowdoin College, Brunswick Maine 04011

## Abstract

In a screen for mutations that alter cell morphology and adhesion as a marker for cell wall composition alterations, an Arabidopsis seedling with root hairs that fail to elongate was identified. Root hairs are rounded and bubble-like, and the mutation was named Bubbles or bbl1. No other visible whole plant phenotypes were observed. Using backcrosses with wild type, and genomic sequencing of pooled F2 individuals with the Bubbles phenotype, 14 candidates for the mutant allele were identified. T-DNA alleles of one of these candidates showed a similar but partial phenotype to bbl1, and indicate that bbl1 is an allele of AT2G39110 previously identified as PBL38 (AvrPphB SUSCEPTIBLE1-LIKE38), a member of receptor kinase subfamily VII. PBL38 has been associated with the response to pathogens. bbl1 is a point mutation that causes a Glutamic acid to Lysine change at position 140, and is complimented by a C-terminal GFP fusion to the coding region of AT2G39110, indicating that the BBL1 gene (PBL38) is involved in correct root hair formation. AT2G39110 is expressed only in roots early in seedling maturation, and the GFP fusion protein localizes to the cell surface, consistent with the prediction that the gene encodes a receptor-like protein kinase. In bbl1 root hairs, the actin cytoskeleton does not form, while in bbl1 roots and other tissues normal actin cytoskeleton is observed.

## Introduction

The cell wall plays an integral role in both maintaining the structure, integrity and adhesion of plant cells. In a screen for mutations that disrupt the cell wall, and specifically cell-cell adhesion (Kohorn et al., 2021) we identified one mutant Arabidopsis that was specifically altered in their root hairs. This screen revealed a number of cell adhesion phenotypes in hypocotyls due to new alleles of genes involved in pectin synthesis (Kohorn et al., 2021; Kohorn et al., 2023). One seedling in the M2 population showed root hairs that failed to elongate, and instead formed small bubbles along the length of the root. The new mutant was called Bubbles or bbl1. Root hairs are formed from a single cell that elongates without further cell division. LRX1 and LXR2 together are required for root hair elongation and double lrx mutants show stunted, often swollen or branched morphologies (Baumberger et al., 2003; Mapar et al., 2021). Numerous other genes have been identified that when mutant, alter root hair morphology (Grierson et al., 2014; Mapar et al., 2021). However, the bubbles phenotype did not exactly match any of these phenotypes. To understand the nature of the bbl1 mutation, and if cell wall components might be affected, bbl1 was identified by genomic sequencing. The result describe a new allele of a previously identified gene, Receptor Kinase Subfamily VII PBL38 (Rao et al., 2018) that had no previous association with root hair morphology.

## Materials and Methods

### Plant growth conditions

Arabidopsis thaliana seeds were sterilized for 5 min in 95% ethanol and then 5 min in 10% bleach and rinsed twice with sterile dH2O. Seeds were then grown in liquid Murashige and Skoog (MS) or MS agar containing media (Sigma-Aldrich, pH 5) with 2% agarose and 1% sucrose or planted directly onto soil. Plated seeds were vernalized at 4°C for 48 h in the dark, moved to the light for 4 h at 20°C and then 5 days in the dark at 20°C. Seeds planted on soil were first vernalized in soli at 4°C for 48 h and grown at 21° C under a 16 h light, 8 hr dark cycle. Plants were imaged using a Nixon D3000 camera.

### Mutant identification

Mutant (M1) plants grown from EMS-mutagenized seeds were grown on soil (Koncz et al., 1992). M2 generation seeds were then collected in 191 pools (each pool contained the progeny of approximately 20-30 plants) and 400 seeds from each pool were then grown for 4 days in the dark in liquid media after sterilization and stained with Ruthenium Red dye (Verger et al., 2016) Liquid media was removed and 3 ml of Ruthenium Red dye (Sigma Corporation, 0.5 mg/ml in dH2O) was applied to seedlings for 2 min in a 10 ml microtiter growth plate. Seedlings were washed twice with 5 ml of dH2O. Hypocotyl staining was then observed under a dissecting microscope, and mutants were transferred to soil.

### DNA extraction and PCR

Three-week old healthy green leaves from plants of interest were collected, frozen in liquid N2, and DNA was extracted as previously described (Kohorn et al., 2016). The indicated genes were PCR amplified according to the manufacturer’s conditions using Titanium Taq DNA polymerase (Takara Bio) using the following primers: Bbl1F <u>5’ GCACGTCCTCTCAGAGACACACAGTAATAATC.</u> Bbl1R<u> 5’</u> CTG AAA CTC CAT GCC TTG ATG AAG ATA AGC G 3’ PCR samples were sequenced by Retrogen.

### elmo1 F2 whole-genome sequencing

The pooled DNA preparations of 100 elmo1 individuals of an elmo1×WT Col0 F2 progeny were sequenced with Illumina genome sequencing technology performed by Novogene. Analysis of the elmo1 F2 allele frequencies was performed using the data file provided by Novogene and the programs artMAP (used to identify the allele frequencies) (Javorka et al., 2019) and IGV used to visualize the genome sequence (Thorvaldsdottir et al., 2013).

### Expression of Bbl-GFP

The coding region of At2g39110 was PCR amplified using the following primers: AT2G39110 Fp 5’ CTC CTT TAC TAG TCA GAT CTA TAG AAG TAA CAA GTT TAT GAG ACC AAG TAG GTA AAT TGG 3’, AT2G39110 R 5’ gacctgcaggcatgcaagctt gagcactatgctctcgtcaaatgaagtaaatagac 3′; The products were cloned into the HindIII BglII restriction enzyme sites of pCambia 1302. Plasmids were then transformed into Agrobacterium and the floral dip method (Clough and Bent, 1998) was used for transformation of bbl1−/− A. thaliana Col0.

### Confocal microscopy

Four-day-old dark grown seedlings were stained for 10 min with 10 μg/mL propidium iodide, and then washed once in dH2O. Hypocotyls were then visualized by confocal microscopy on a Leica SP8 microscope using a 10× objective, a 514 nm excitation laser and an emission spectrum of 617 nm. A z-stack was then created for the seedling using the Leica SP8 software. GFP fusion proteins were detected by scanning on the Leica SP8 using excitation 488 nm, emission 510-515 nm.

## Results and Discussion

The bubble mutant was identified using ruthenium red staining of dark grown hypocotyls. Ruthenium red binds to de-esterified pectin, and can penetrate roots, root hairs, but not hypocotyl cell walls, and highlights tissue morphology so that any changes are more easily detected on a dissecting microscope (Sola et al., 2019; Verger et al., 2016) . To obtain this mutant, 5000 wild type (WT) Arabidopsis seeds were mutagenized with ethyl methanesulfonate (EMS), self-crossed and the M2 (mutagenized F2) population were collected in 191 pools (Kohorn et al., 2021). Fig 1 shows the ruthenium red stained seedlings for bbl1 and wild type (WT), where only the roots are affected. Seedlings were also stained with propidium iodide and visualized on the confocal microscope, thus providing a more clear image of the mutant phenotype relative to wild type (Fig 2).

**Fig 1.**
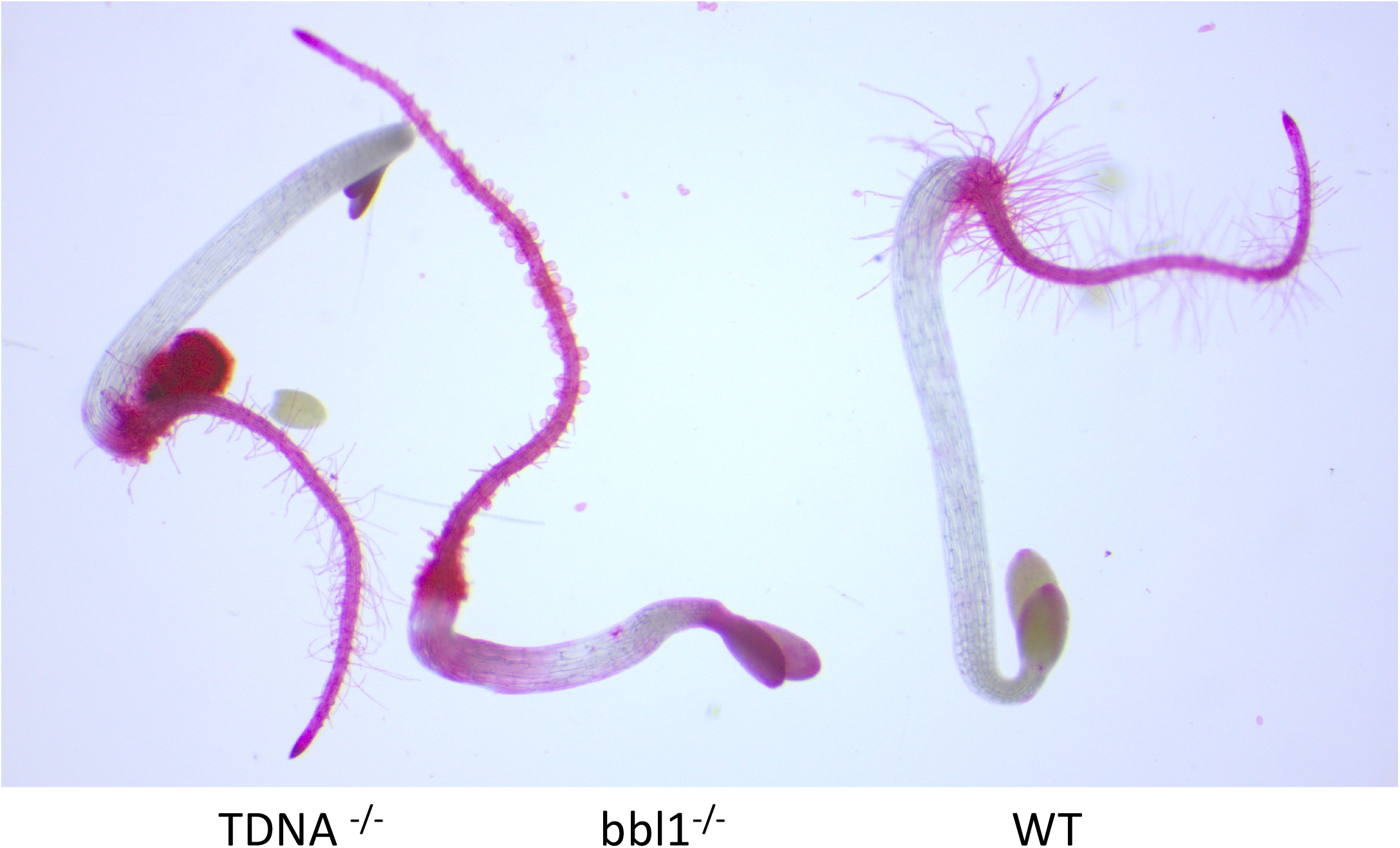
Ruthenium Red stained dark grown hypocotyls of the indicated genotype.

**Fig 2.**
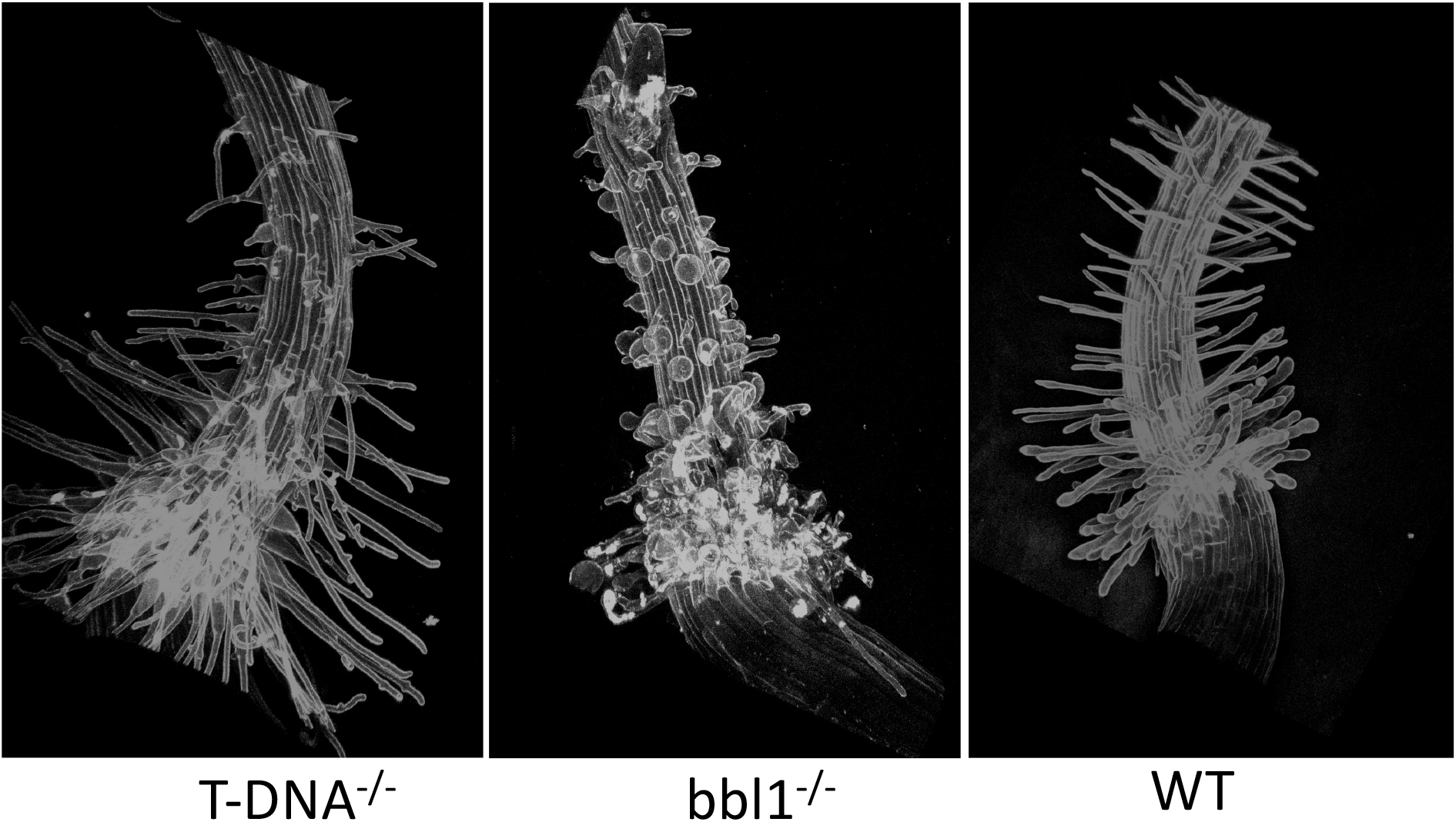
Prodium Iodide stained confocal images of dark grown hypocotyls of the indicated genotype.

To identify the mutation causing the bbl1 phenotype, bbl1 was crossed to WT, and the F1 seeds were first grown in liquid in the dark for 5 days to observe the phenotype. All F1 appeared wild type, indicating that the bbl1 allele is recessive. F1 seeds were also sown in soil and the resulting plants allowed to self-cross. Of the F2 offspring, a quarter showed the bubble phenotype in dark grown hypocotyls, as expected. In the F2 one quarter were expected to be homozygous for the bbl1 mutation, but other EMS-induced background mutations present in these F2 bbl1 −/− would be expected to have segregated and appear at less than 100% frequency. Mutations that occur with 100% frequency in this F2 population were identified by whole-genome sequencing of 400 pooled F2 seedlings with the bbl1 phenotype. Allele frequencies were identified using artMAP software (Javorka et al., 2019) and 14 mutations occurring with 100% frequency were identified (Table 1). However, only four mutations in genes (AT2G35350, AT2G37035, AT2G38670, AT2G39110) resulted in an amino acid change and were more likely to be the causative allele. One other contained a mutation in the 5’ untranslated region and might affect expression (AT2G39180). Several other mutations were in mitochondrial genes, and were therefore not pursued as the inheritance of the allele was not consistent with the mitochondrial segregation. The poltergeist allele was also not pursued since the phenotype of mutations in this gene do not match that of bbl1 (Song et al., 2020).

**Table 1.**
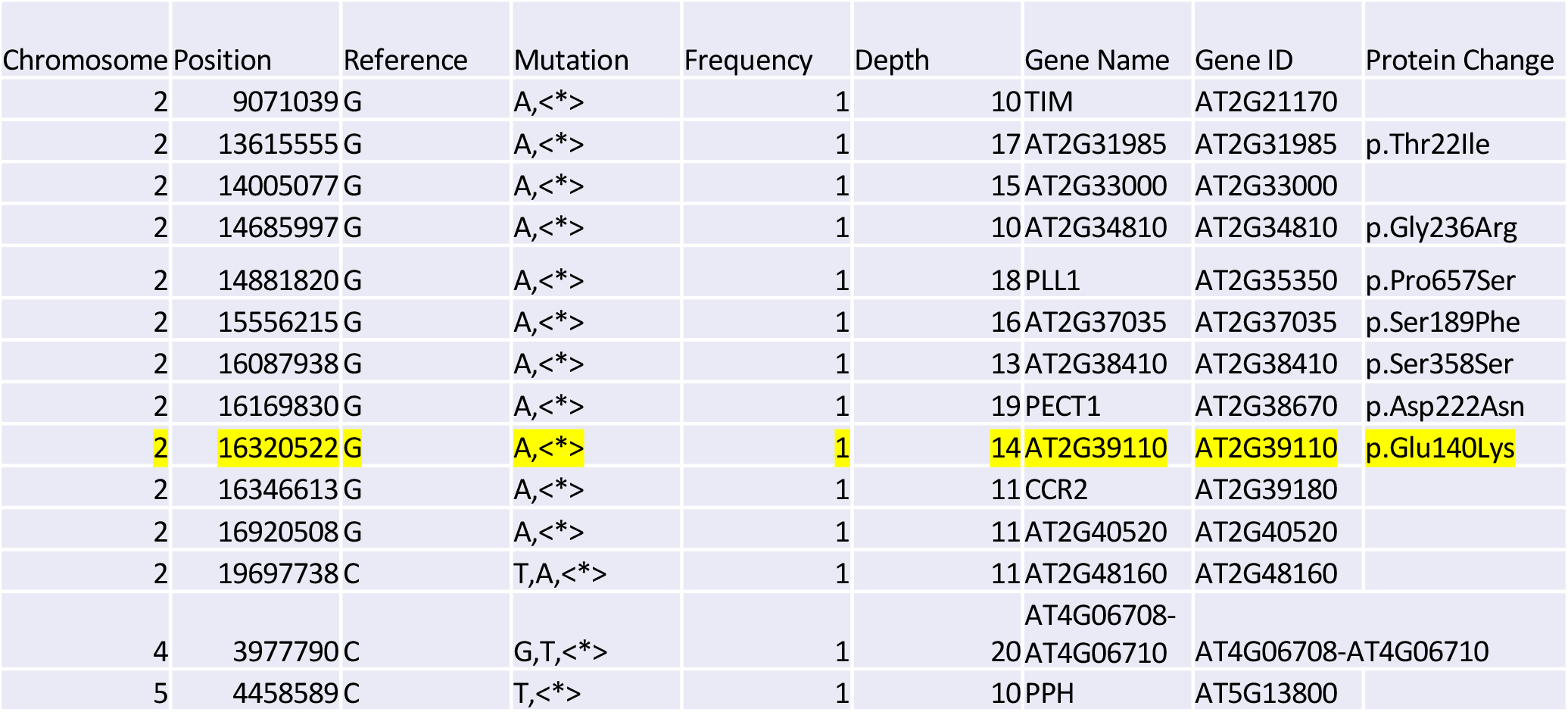
List of alleles detected at 100 % frequency in the F2 segregating population of bbl1 crossed with WT Arabidopsis. Table of allele frequency of pooled F2 bbl1 x WT

To determine which of the 5 loci was affected in bbl1, plants having available T-DNA insertions in were ordered from the Arabidopsis stock center (Arabidopsis Biological Resource Center) and dark grown seedlings homozygous for the T-DNA insertion were stained with ruthenium red. Only one insertion line showed a root hair phenotype of any kind, and this was in AT2G39110, Sail 416 A03 having an insertion in the second intron (Fig 1 and Fig 2, Fig 3). Interestingly in the TDNA line, while many root hairs especially at the hypocotyl junction formed bubbles, many along the root appeared normal or had lateral bulges or branches. It is likely that since the T-DNA insertion is in an intron this allele has less of an effect than does the single point mutation of bbl1 that substitutes a Lysine for a Glutamic acid within the putative kinase domain (Fig 3). AT2G39110 was identified previously as PBL38 (AvrPphB SUSCEPTIBLE1-LIKE38), a member of receptor kinase subfamily VII (Rao et al., 2018). PBL38 has been associated with the response to pathogens, is subject to ubiquitylation and regulated turnover, but little else is known of this receptor-like kinase (Bai et al., 2022). AT2G39110 is expressed only in roots early in seedling maturation (The Arabidopsis Information resource TAIR), and while root hairs appear abnormal, the remaining root structures are indistinguishable from WT. Hence it is not unexpected that a mutation in the BBL1 locus would only provide a root phenotype.

**Fig 3.**
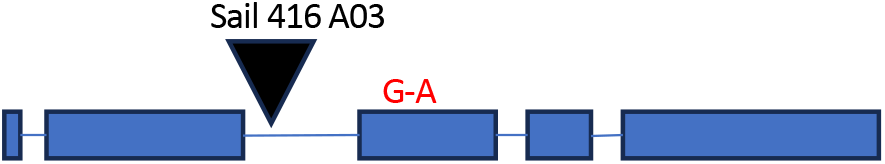
Cartoon of the exons (blue box) introns (blue line) of the BBl1 locus, indicating the location of the T-DNA insertion (black arrow) and the point mutation at codon 140 (red G-A).

To verify that bbl1 is the causative allele for the bubble phenotype, a gene for a carboxyl-terminal fusion to the coding region of AT2G39110, driven by 1.5 kb of the native promoter was transformed into the bbl1 ^-/-^ plant, and the T1 were scored for phenotype and GFP expression. Roots of T2 individuals are shown in Fig 4, and expression of GFP was only detected in roots (Fig 4B) and root hairs (Fig 4 A and no other tissues of the plants, consistent with the reported activity of the gene. All plants expressing GFP were wild type, indicating that the BBL1-GFP fusion compliments the bbl1 mutant, and that *BBL1* gene (PBL38) is involved in correct root hair formation. GFP fluorescence is detected only at the cell surface, and not in internal organelles or the cytoplasm, consistent with the prediction that PBL38 is a receptor-like kinase.

**Fig 4.**
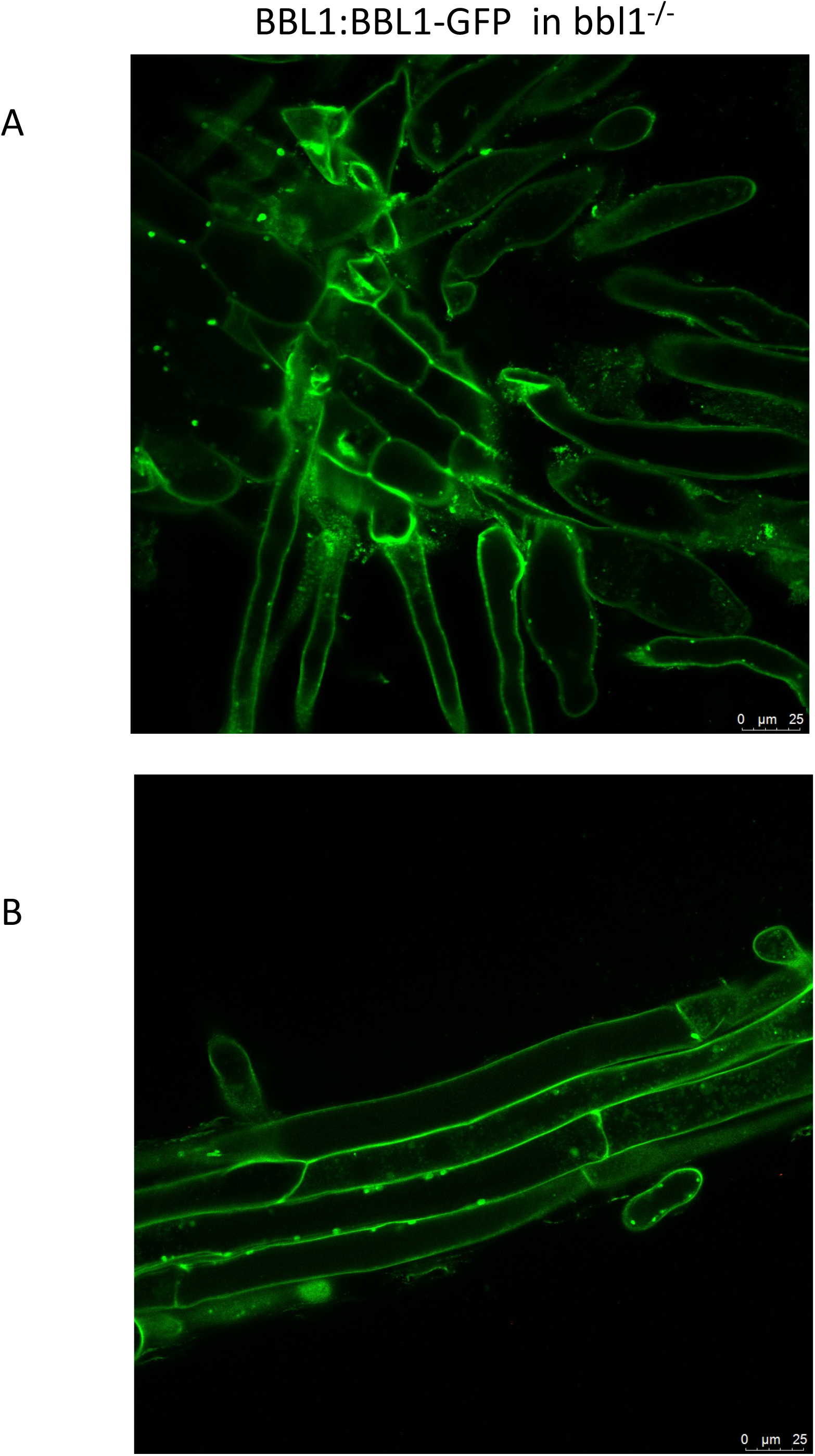
Confocal image of FPF fluorescence from root/hypocotyl junction (A) and the root (B) for bbl1 transformed with BBL1 promoter driven BBL1-GFP fusion protein. Bar indicates 25 um.

To gain an understanding of what genes might be downstream of a potential activation of the BBL1 receptor, total RNA was isolated from WT and bbl1 ^-/-^ dark grown hypocotyls, and RNA seq analysis was performed. While there were some genes that were up or down regulated in bbl1 ^-/-^ plants, relative to WT, there appeared to be no consistent pattern or changes in gene expression. It was hoped that perhaps BBL1 target genes might be identified, but this was not evident from the analysis. However, a more accurate data set needs to be generated from roots, and perhaps root hairs as small differences in root hairs may have been diluted by hypocotyl and root tissues that do not differ between bbl1 and WT.

To determine why the root hairs might be forming bubbles and not elongating, the bbl1 allele was crossed with a plant expressing a Fimbrin-GFP in order to visualize the actin cytoskeleton (Vaskebova et al., 2018). Plants homozygous for the bbl1 allele were screened for in the F2 population of a cross between Fimbrin-GFP and bbl1 ^-/-^ , using PCR and primers of BBL1 and sequencing of the product. Fig 5 shows the Fimbrin-GFP fluorescence from bbl1 ^-/-^ seedlings compared to WT also expressing Fimbrin-GFP. WT root hairs have visible arrays of actin cytoskeleton, yet the bubble-like root hairs have no organized filaments, and instead a faint background of fluorescence perhaps representing a pool of unpolymerized actin. Actin filaments are detected in the bbl1 ^-/-^ roots, which correspondingly do not show an obvious bbl1 phenotype.

**Fig 5.**
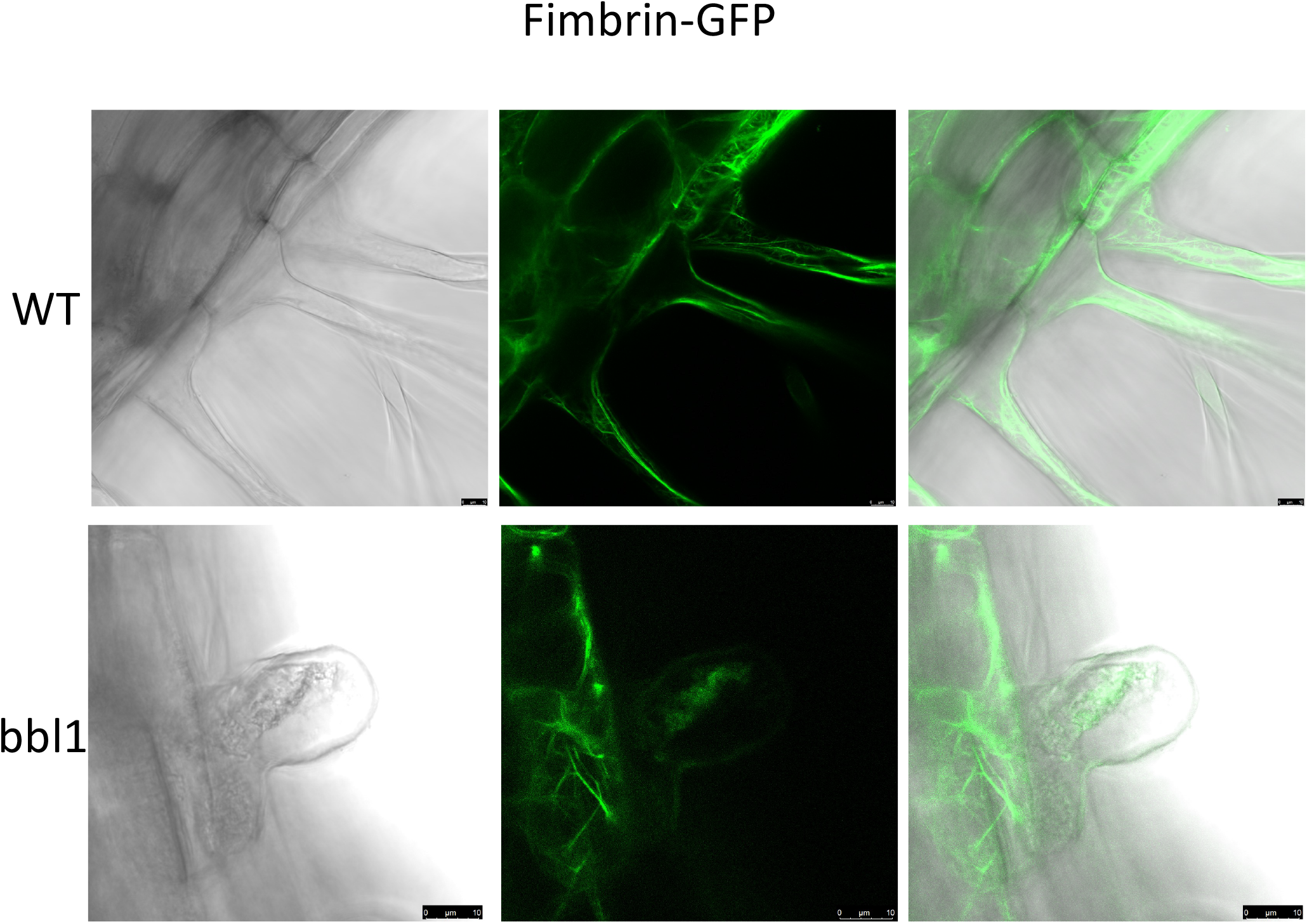
Expression of Fimbrin-GFP fusion protein to detect actin in WT and bbl1^-/-^ . Left panel is white light image, center panel is GFP fluorescence, right images are overlay of the two left panels. Bar indicates 10 um. Images were taken on a confocal microscope.

In conclusion, we provide evidence that alteration of a single amino acid in BBL1 (PLB38), a receptor kinase of subfamily VII, leads to bubble-like root hairs. No other phenotypes were detected as the gross morphological level, nor were there any changes in gene expression patterns that suggested potential targets for this putative receptor kinase. A dramatic loss of an organized actin cytoskeleton in the bubble-like root hairs was detected in bbl1 ^-/-^, suggesting that BBL1 (PLB38) either directly, or indirectly regulates actin assembly or stability.

## Acknowledgements

The work was supported by Linnean Chair Funds from Bowdoin College. Thank you to Sue Kohorn, Isabell Ball, Garrison Asper, Frnacis Zorensky and Jacob Dexter-Meldrum (in memory).

## Notes

### Competing Interest Statement

The authors have declared no competing interest.

